# A sporulation-independent way of life for *Bacillus thuringiensis* in the late stages of an infection

**DOI:** 10.1101/2022.05.09.491210

**Authors:** Hasna Toukabri, Didier Lereclus, Leyla Slamti

## Abstract

The formation of endospores has been considered as the unique mode of survival and transmission of sporulating Firmicutes due to the exceptional resistance and persistence of this bacterial form. However, the persistence of non-sporulated bacteria (Spo^-^) was reported during infection in *Bacillus thuringiensis*, an entomopathogenic sporulating Gram-positive bacterium. In this study, we investigated the behavior of a bacterial population in the late stages of an infection as well as the characteristics of the Spo^-^ bacteria in the *B. thuringiensis*/*Galleria mellonella* infection model. Using fluorescent reporters coupled to flow cytometry as well as molecular markers, we demonstrated that the Spo^-^ cells constitute about half of the population two weeks post-infection (pi) and that these bacteria present vitality signs. However, a protein synthesis and a growth recovery assay indicated that they are in a metabolically slowed-down state. Interestingly, they were extremely resistant to the cadaver environment which proved deadly for *in vitro*-grown vegetative cells and, strikingly, did not support spore germination. A transcriptomic analysis of this subpopulation at 7 days pi revealed a signature profile of this state. The expression analysis of individual genes at the cell level suggests that iron homeostasis is important at all stages of the infection, whereas the oxidative stress response seems of particular importance as the survival time increases. Altogether, these data show that non-sporulated bacteria are able to survive for a prolonged period of time and indicate that they engage in a profound adaptation process that leads to their persistence in the host cadaver.

## INTRODUCTION

The ability of bacteria to face environmental fluctuations is essential to their survival. These stresses include nutrient limitation or depletion, variations of pH, temperature, oxygen level, osmolarity, radiation and chemical exposure. In conditions that do not support their growth, some Firmicutes, such as *Bacillus* species, have evolved the ultimate mode of resistance, sporulation [1]. This survival mode requires the activation of a specific and complex developmental pathway that leads to the formation of a spore, a highly resistant dormant cell that allows these species to survive in extreme conditions during extended periods [2–5].

Due to their resistant nature, the formation of endospores was considered the unique mode of survival and transmission of these sporulating species. However, it was recently shown that a *Bacillus subtilis* sporulation mutant was able to survive deep starvation during several months by entering an oligotrophic state that supports extreme slow growth [6]. Moreover, spontaneous mutants in *spo0A*, the gene specifying the master regulator of sporulation [7, 8], were isolated alongside spores after a month-long incubation of the food-borne pathogen *Bacillus cereus* under abiotic conditions in groundwater [9]. Furthermore, a *B. cereus* strain impaired in sporulation was shown to persist for several days on ready-to-eat vegetables and retained its ability to cause illness in a mammalian host [10]. In addition, it was shown that spores represented only 30% of the bacterial population that survived in an insect cadaver 4 days pi (dpi) with *Bacillus thuringiensis* and that a sporulation mutant was able to persist in the cadaver during that time [11]. *B. thuringiensis* is an entomopathogenic bacterium that belongs to the *B. cereus sensu lato* group comprising several species that are pathogenic to humans and animals, notably *B. cereus sensu stricto*, which is responsible for foodborne toxi-infections as well as systemic infections, and *Bacillus anthracis*, the agent of anthrax [12]. *B. thuringiensis* was shown to carry out a full infectious cycle in the *Galleria mellonella* insect larvae model. This process is composed of three major phases controlled by quorum-sensing systems [13]: virulence that allows the bacteria to invade and kill the host [14], necrotrophism which permits the bacteria to feed on the cadaver [11], and sporulation. These processes were reported in insects using fluorescent reporters to detect the activity of the regulators responsible for these states at the cell level [15]. Sporulation was shown to only occur in a part of the subpopulation that had activated the necrotrophic regulon. This is due to the dual nature of the NprR regulator that acts as an activator of the necrotrophic regulon when bound to its cognate signaling peptide NprX, and as a phosphatase negatively regulating sporulation in its apo-form [16]. Necrotrophism was reported to be activated during the whole experiment time-span (*i.e.* 4 days) in bacteria that did not engage in sporulation. A category of cells, which did not express the necrotrophic or the sporulation reporters in the host, was identified and designated Nec^-^/Spo [15, 17]. Most of these bacteria were identified as viable using a cell death marker at 3 dpi and the results suggested that they were not in exponential or stationary phase [18], (and unpublished).

In this study, we investigated the composition of the bacterial population during the late stages of an infection as well as the characteristics and the proportion of the non-sporulated bacterial population in the *B. thuringiensis*/*G. mellonella* infection model. We used an unstable fluorescent reporter coupled to flow cytometry to determine the actual duration of necrotrophism. We also examined the metabolic state of the Spo^-^ cells using protein synthesis and growth recovery assays on rich or insect-based medium. We performed a transcriptomic analysis of this subpopulation at 7 dpi that revealed a signature profile of this state.

## MATERIALS AND METHODS

### Bacterial strains and growth conditions

The acrystalliferous *Bacillus thuringiensis* 407 Cry^−^ strain (*B. thuringiensis* 407^-^) [19] was used as the parental strain to create all the strains used in this study. *Escherichia coli* strain DH5α [20] was used as the host strain for plasmid construction. *E. coli* strain ET12567 [21] was used to prepare DNA prior to electroporation in *B. thuringiensis*. Cells were grown under shaking at 30°C or 37°C in LB medium (1% tryptone, 0.5% yeast extract, 1% NaCl) or HCT medium (0.7% casein hydrolysate, 0.5% tryptone, 0.68% KH_2_PO_4_, 0.012% MgSO4·7H2O, 0.00022% MnSO4·4H2O, 0.0014% ZnSO4·7H2O, 0.008% ferric ammonium citrate, 0.018% CaCl2·4H2O, pH 7.2). The bacteria were stored at −80°C in LB containing 15% glycerol. When required, the medium was supplemented with antibiotics at the following concentrations: 100 µg/mL ampicillin (for *E. coli*), 200 µg/mL kanamycin, 10 µg/mL erythromycin (for *B. thuringiensis*).

### Plasmids and strains construction

DNA manipulations are detailed in the supplemental material. All the plasmids, strains and oligonucleotide primers used in this study are listed in Tables S1, S2 and S3 in the supplemental material.

### *Infection of* Galleria mellonella

Intrahemocoelic injection experiments with *Galleria mellonella* were carried out as described previously [14]. Briefly, last instar larvae were infected by intrahaemocœlic injections with 2.10^4^ vegetative cells in exponential phase (OD_600_=1). Inocula were numerated after plating onto LB agar medium. Infected larvae were incubated at 30°C under a Nikon CoolPix P1 on time-lapse photography mode and pictures were taken every 10 min to determine the time of death for each larva. Larvae that died within the same hour were used for each time point. A larva was considered dead at the time its movements stopped if followed by melanization (see supplemental material for details, fig S1).

### *Extraction of* B. thuringiensis *from* G. mellonella *cadavers*

Isolation of bacteria from insect cadavers was adapted from Verplaetse et al., 2015, [15]. Cadavers were crushed with a 1 mL tip in a 1.5 mL Eppendorf tube. 1 mL of saline was added and the tube was vortexed. 750 µL of the suspension were transferred to a new 1.5 mL tube and centrifuged for 45 sec at 13000 rpm. The pellet was resuspended in 750 µL of saline and loaded in a 1 mL syringe fitted with a cotton filter in order to isolate bacterial cells from cadaver debris. The filtrate was then centrifuged for 30 sec at 13000 rpm, fixed in 4% formaldehyde in saline for 15 min, washed in saline, resuspended in GTE buffer [22] and kept at 4°C until flow cytometric or microscopy analysis.

### Flow cytometric analysis

Bacteria were diluted in filtered saline and fluorescent events were measured on a CyFlow Space cytometer (Sysmex Partec, France). Specifications of the apparatus and flow cytometric analyses are described in the supplemental material.

### Microscopy

Bacteria were observed with an AxioObserver.Z1 Zeiss inverted fluorescence microscope with a Zeiss AxioCam MRm digital camera and with Zeiss fluorescence filters. GFP, 5(6)-CFDA, DiBAC4(3), Sytox Green were imaged using the 38 HE filter (excitation: BP 470/40, beam splitter: FT 495, emission: 525/50) and mCherry was imaged using the 45 HE filter (excitation: BP 590/20, beam splitter: FT 605, emission: 620/14). Images were processed using the ZEN software package (Zeiss).

### Sporulation assay

The sporulated population was assessed by plating and by flow cytometry. Each larva cadaver was ground in 3 mL of saline on ice using an T10 Ultra-Turaxx apparatus. The volume was then adjusted to 10 mL. To determine the number of total cells and the number of heat-resistant spores, the bacteria were enumerated by plating on LB agar plates following serial dilutions before and after heat treatment at 80°C for 12 min. For flow cytometry, fixed bacteria (as described above) harboring the red fluorescent sporulation reporter *PspoIIQ-mCherry* were analyzed. Red fluorescent events were counted among the total cell count to calculate the percentage of sporulation.

### Induction of gfp expression

Bacteria were extracted from insect cadavers as described above or were harvested from HCT cultures at 30°C at OD_600_=1 (exponential phase of growth) and OD_600_=6 (stationary phase of growth) *via* centrifugation at 13000 rpm for 30 sec, washed once in saline and resuspended in conditioned HCT medium (obtained after filtration of a 10 h culture of *B. thuringiensis* 407^-^ in HCT) to preserve bacteria while preventing cell growth. Half of the suspension was supplemented with xylose at 25 mM final concentration. For each sample, an aliquot was fixed as described above at the time of xylose addition and after 1 and 2 h of incubation with shaking at 30°C. The samples were kept at 4°C until flow cytometric or microscopy analysis.

### Growth recovery assessment of the bacteria extracted from insect cadavers

Culturability was assessed by plating the bacteria on two different media, LB agar and Insect agar, and by time-lapse microscopy. The bacteria were extracted from insect cadavers as described above or were harvested from *in vitro* cultures in LB at 30°C at OD_600_=1 (exponential phase), OD_600_=8 (stationary phase) and from a 48 h culture (composed of about 65% of spores) *via* centrifugation at 13000 rpm for 30 sec, washed once in saline, serial diluted and plated onto LB plates containing erythromycin. 16 hours after incubation at room temperature, colonies were photographed with a Canon Eos 750D under a binocular stereo microscope Leica MZFLIII. Colony area was determined using the Image J software [23].

To prepare Insect agar plates, 12 larvae cadavers were harvested at 7 dpi and crushed in 50 mL saline then filtered as described above. Agar prepared in saline was then added at a final concentration of 1.5%. The bacteria extracted from insect larvae or harvested from LB cultures, in the same manner as above, were serial diluted and plated onto LB plates supplemented with erythromycin and Insect agar plates supplemented with erythromycin. Colonies were counted 16 hours after incubation at room temperature for LB plates and after 4 days for Insect agar plates.

For time lapse microscopy, medium pads were prepared with LB supplemented with 1% agarose to fill a 125-µl GeneFrame (Thermo Fisher Scientific, Waltham, MA, U.S.A.). After polymerization, 2-mm-wide strips were cut and three strips were placed in a new gene frame. Bacteria were extracted from insect larvae or harvested from LB cultures as described above and washed in saline. 2 µL of cells were then pipetted onto the strips before the frame was sealed with a coverslip. *B. thuringiensis* development was monitored at 30°C in a temperature-controlled chamber with the microscope described above. Phase-contrast and fluorescence images were taken every hour for 5 h. Only bacteria extracted from insect cadavers at 0 h were assigned to a category (elongation, division, lysis, inactivity). Newly-formed daughter cells were excluded from the analysis.

### Molecular dye staining

Bacterial cells were harvested from LB cultures or insect cadavers as described above and resuspended in 1 mL of saline. Samples were diluted to 10^7^ cells/mL in filtered saline supplemented or not with one of the dyes. 5(6)-CFDA (Biotium, Fremont, CA, USA), DiBAC4(3) (Sigma-Aldrich, Saint-Louis, MO, USA) and Sytox Green (Invitrogen, Eugene, OR, USA) were suspended in DMSO and used at a final concentration of 5 μM (5(6)-CFDA) or 0.5 μM (DiBAC4(3) and Sytox Green). Cells were incubated at room temperature for 15 min in the dark. Stained samples were washed once in saline before flow cytometric or microscopy analysis. Unstained and heat-shocked cells at 90°C for 15 min were used as control.

### RNA-Seq analysis

RNA was extracted from shaken liquid cultures and from dead larvae at 7 dpi as described in the supplemental material. Nanodrop and Bioanalyzer instruments were used for quantity and quality controls (6,8 ≤ RINs ≤ 10). Disrupting cells broke Spo^-^ bacteria and left spores intact. Moreover, extraction of RNA from spore preparations using our protocol resulted in a very low yield of poor quality. However, we can not exclude a minute contamination with RNA from spores.

Library preparation including ribosomal RNA depletion (RiboZero) and sequencing was performed by the I2BC platform (Gif-sur-Yvette, France). Sequencing was conducted on an Illumina NextSeq machine using NextSeq 550/500 Mid Output Kit v2 to generate paired end reads (2×75).

Data processing was performed by the I2BC platform including the following steps: demultiplexing (with bcl2fastq2-2.18.12), adapter trimming (Cutadapt 1.15), quality control (FastQC v0.11.5), mapping (BWA v0.6.2-r126) against *B. thuringiensis* 407 strain. This generated between 9 M and 18 M of uniquely mapped reads per sample and counts for 6530 genes. Differentially expressed genes (DEG) were determined using the R package “DESeq2” (v1.30.1) to estimate p-values and log2 fold-changes. DEG with padj ≤ 10^-2^ and log2FC≤-2 or log2FC ≥ 2 were used for further analyses. Gene expression data are being deposited at the public repository Gene Expression Omnibus.

Using the functional re-annotation database available in the laboratory (see details in supplemental materials) of the *B. thuringiensis* 407 genome (chromosome: INSDC CP003889.1, plasmids: INSDC CP003890.1 to CP003898.1), GO terms, COG letters and *B. subtilis* homologous genes were assigned to our data-set with Microsoft SQL server 2019. Venn Diagrams were constructed with Venny 2.1.0 [24], 2007-2015 https://bioinfogp.cnb.csic.es/tools/venny/index.html). Using Subtiwiki [25], KEGG orthology database [26] and COG letters, we retrieved genes from these functional categories: oxidative stress, iron homeostasis, DNA recombination and repair, necrotrophism, sporulation, and germination (see supplemental material, table S4, S5, S6, for necrotrophism, sporulation and germination, respectively). Histograms and heatmaps of DEG according to these functional categories were then constructed using Prism 8 software (GraphPad).

### Statistical analysis

The data were analyzed using the Prism 8 software (GraphPad).

## RESULTS

### Non-sporulated bacteria survive in insect cadavers for at least 14 days post-infection

A previous study recorded the sporulation rate of *B. thuringiensis* in insect larvae cadavers and showed that heat-resistant spores were detected 24 hpi and reached a plateau at 30% of the bacterial population from 48 hpi until the end of the experiment 4 dpi [11]. This result indicated that a large part of the bacterial population was able to survive for 4 days in these conditions without resorting to sporulation. In order to determine if the non-sporulating cells persisted during a longer period of time, and if the sporulation rate remained constant, we monitored the fate of the bacterial population during 14 days using two complementary methods. *B. thuringiensis* cells harboring P*spoIIQ’mcherry*, a transcriptional fusion between the promoter of *spoIIQ*, a sporulation gene activated when the cells are committed to sporulation [27], and the *B. thuringiensis*-optimized *mcherry* fluorescent reporter gene [15], were injected into *G. mellonella* larvae. Sporulation was then assayed by plating and by flow cytometry (as described in the Materials and Methods section). In our conditions, flow cytometry analysis of P*spoIIQ’mCherry* cannot discriminate between bacteria irreversibly engaged in sporulation and sporulated cells as the mCherry protein remains fluorescent in spores. In the plating assay, we cannot exclude that some spores will not be able to germinate and form colonies on LB agar plates. These two methods were therefore implemented to give the closest assessement of the sporulated and non-sporulated subpopulations. Larvae died between 8 to 12 hpi (fig S1) and bacteria were extracted from insect cadavers from 16 hpi to 14 dpi for analysis. The plating assay showed that the total number of CFU per larva remained stable during 14 dpi (fig S2). mCherry-positive bacteria (Fig 1.a and 2.a) and heat-resistant spores (Fig. S2) were detected starting 1 dpi and represented about 40 and 50 % of the total population, respectively. Accordingly, microscopy results indicate that the mCherry-positive events mainly correspond to spores from the third day post-infection onwards (fig S2 and data not shown). The sporulation rate remained stable during 14 days and was comprised between 30% and 55% indicating that up to 70% of the cells could be in a non-sporulated form. We cannot exclude that germination occurred in insect cadavers as a few spores had lost their refringence and remained fluorescent when observed by microscopy (data not shown). However less than 5% of the bacteria presented this profile (data not shown). Comparison of the data using a Bland-Altman plot (fig 2.b) showed that the results obtained using both methods are similar. Indeed, the bias value is 3.2 and the SD of bias 7.3, indicating that there is less than 10% difference in the sporulation percentage when comparing plating and flow cytometry. This indicates that both methods are equivalent for evaluating sporulation in these conditions. Taken together, these results show that sporulation is not the only survival strategy in insect cadavers even in late-infection stages.

**Figure 1.**
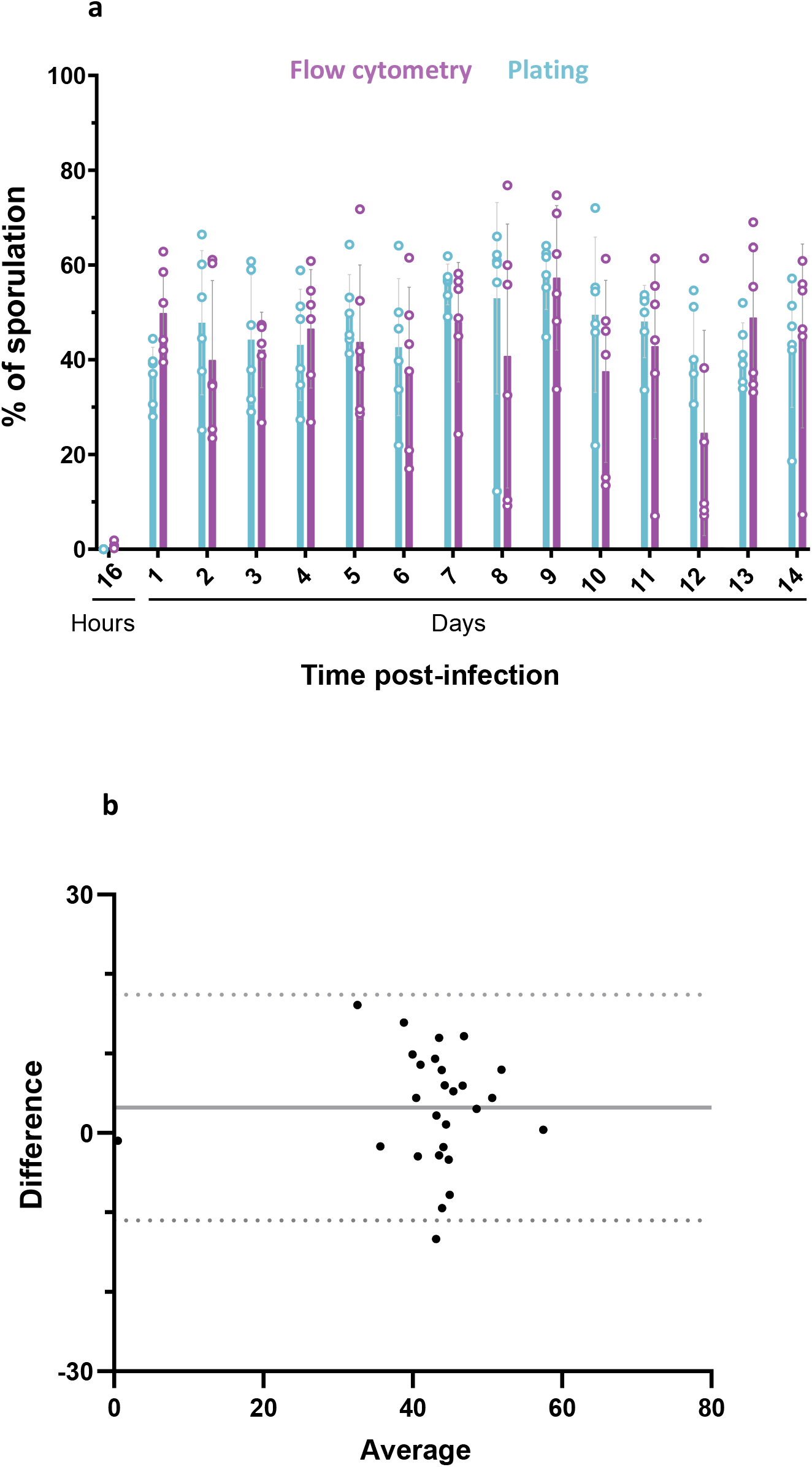
Monitoring of the sporulated subpopulation during long-term infection. **a.** The percentage of sporulation was determined daily for 14 days after intrahemocoelic infection of *G. mellonella* with Bt (pP*nprA’gfp_Bte_AAV-*P*spoIIQ’mCherry*) by plating onto LB agar (blue) or by flow cytometry (purple). 6 larvae were crushed at the time points indicated and serial dilutions of the homogenate were directly plated onto LB agar for total population numeration, or after heating at 80°C during 12 minutes to account for heat-resistant spores. For the flow cytometry assay, the same samples were fixed and red fluorescent events were discriminated in cytograms (as described in the Materials and Methods) and counted as sporulating bacteria. The total population was determined by the number of total events. Each symbol represents the data relative to bacteria extracted from one larva. The data are the result of two independent experiments and the error bars show the standard deviation from the mean. **b.** Bland-Altman comparison plot between flow cytometry and plating sporulation assessment. The difference between plating and flow cytometry is plotted on the Y axis and the average of the values obtained by both method is plated on the X axis. The black line represents the bias value (3,198) and dotted lines indicates the 95% limits of agreement. Data were collected from 27 larvae.

**Figure 2.**
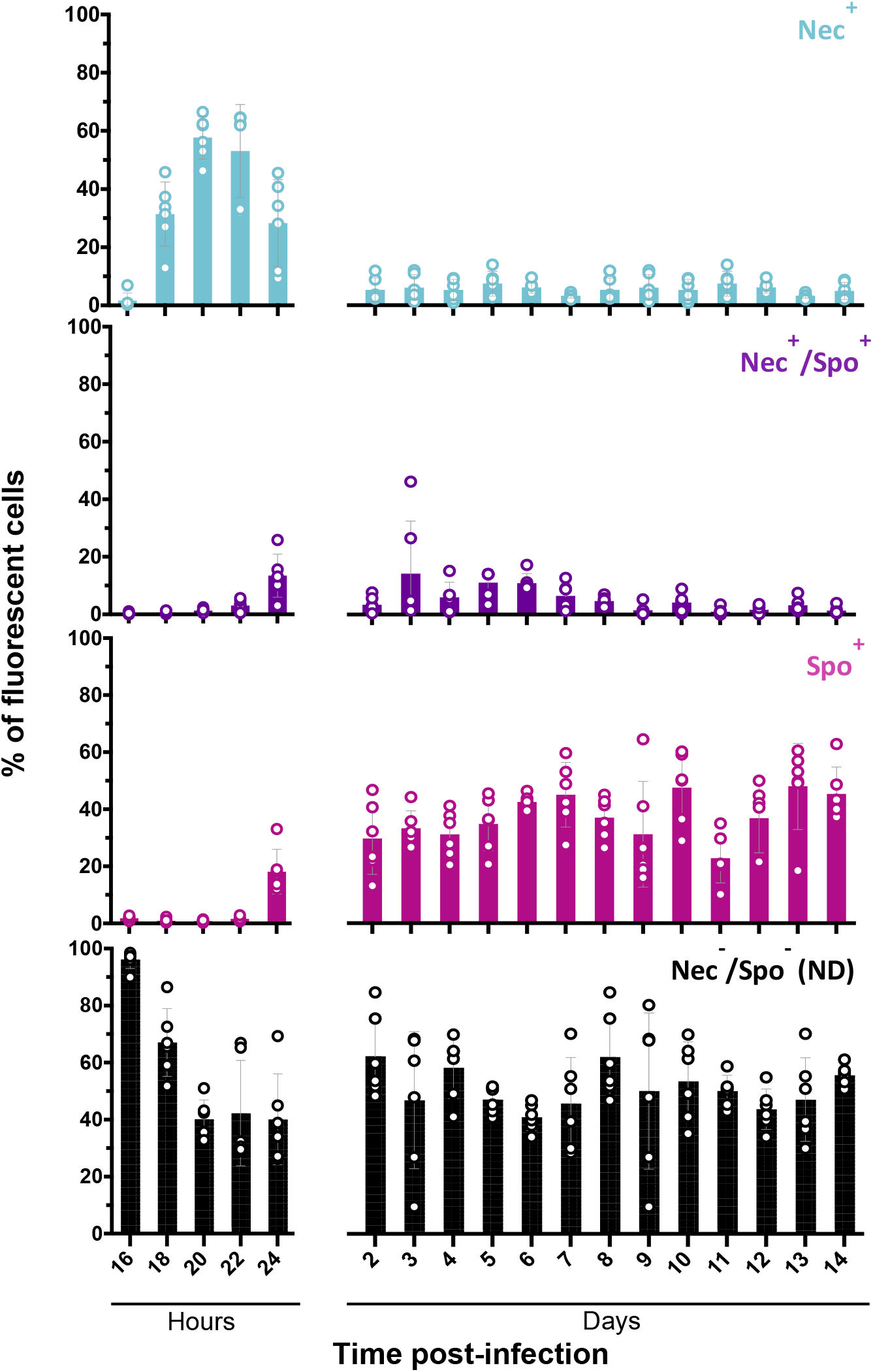
Necrotrophism and sporulation promoter activities in bacterial cells during long-term infection. Flow cytometry analysis of Bt (pP*nprA’gfp_Bte_AAV-*P*spoIIQ’mCherry*) cells during *G. mellonella* infection. Cells were extracted from 6 larvae cadavers at the time points indicated during 14 days after intrahemocoelic infection. The percentage of the total cell count represented by each population was discriminated in cytograms and is presented as a function of time (as described in the Materials and Methods section). Each population phenotype is associated with a color: Nec^+^ in blue for cells expressing the necrotrophic marker, Spo^+^ in pink for cells expressing the sporulation marker, Nec^+^/Spo^+^ in purple for cells expressing both markers, ND in black for cells that do not express any of the markers used. Each symbol represents bacteria extracted from one larva. The data are the result of two independent experiments and the error bars show the standard deviation from the mean.

### Necrotrophism is a transient state occurring at early stages of survival in insect cadavers

To assess the proportion of bacteria in the necrotrophic state during the late stages of infection in insect larvae, we monitored this state for 14 dpi using a transcriptional fusion between the promoter of *nprA*, an NprR-regulated gene [28], and the reporter gene *gfp_Bte_AAV*, encoding an unstable Gfp [18]. This reporter gene was previously used to detect vegetative cells *in vivo* and showed its relevance as a molecular tool to assess transient gene expression during infection [18]. We associated this construct to the P*spoIIQ’mcherry* fusion. This combination of reporters allowed us to determine the fraction of the population that was able to persist in the late stages of infection without entering sporulation or activating the necrotrophism pathway.

*G. mellonella* larvae were infected by intrahemocoelic injection with the Bt (pP*nprA’gfp_Bte_AAV*-P*spoIIQ’mCherry*) strain. Figure 2 shows that expression of the P*nprA’gfp_Bte_AAV* fusion started between 16 and 18 hpi with about 30% of the bacteria in the necrotrophic state (Nec^+^) at 18 hpi. The number of Nec^+^ bacteria peaked at 20 hpi to near 60% and decreased to around 30% at 24 hpi. Expression of the P*spoIIQ’mCherry* fusion started between 22 and 24 hpi with about 20% of the bacteria expressing the sporulation reporter only (Spo^+^) and 15% of the population expressing both the necrotrophic and sporulation reporter (Nec^+^/Spo^+^). At 2 dpi the total number of Nec^+^ and Nec^+^/Spo^+^ bacteria decreased to about 10% of the population and remained at that level until the end of the experiment. These data show that necrotrophism is a transient state in which the bacteria remain during a limited period after the death of their host. Furthermore, our results reveal that the previously reported proportion of Nec^-^/Spo^-^ cells was underestimated.

### The majority of the non-sporulated late-infection stage bacteria present cellular vitality

We analyzed the behavior and features of the Spo^-^ bacteria during infection using molecular dyes to assess bacterial vitality. We used three commercial dyes to examine different vitality features of these bacteria: enzymatic activity by assessing esterase activity with 5(6)-CFDA (fig 3.a), membrane potential with DiBAC4(3) (fig 3.b) and membrane integrity with Sytox Green (fig 3.c) [29]. Sytox Green staining was previously successfully used on cells extracted from insect cadavers to detect mortality among the 3 dpi non-necrotrophic cells [18]. Strain Bt (pP*spoIIQ’mCherry*) cultured in LB and harvested in exponential growth phase (Expo) and stationary growth phase (Stat), or extracted from insect cadavers at 1, 3 and 7 dpi, was incubated with the molecular dyes listed above. Cells were then analyzed by flow cytometry (fig 3) and microscopy (fig S4) to quantify the stained Spo^-^ cells. The results show that 5(6)-CFDA stained 90% of Expo and Stat cells, and 67% to 72% of insect cadavers cells (fig 3). However, this difference was not statistically significant. Less than 16% of Expo and Stat cells and approximately 30% of insect cadavers cells were marked with DiBAC4(3) and Sytox Green (fig 3.b and 3.c). Bacteria extracted from insect cadavers presented a more heterogeneous profile as the percentage of marked cells ranged from 15% to 60% of the Spo^-^ bacteria. Our results indicate that a portion of the Spo^-^ cells extracted from insect cadavers was damaged, but the majority of these bacteria presented esterase activity, membrane polarization and membrane integrity, indicating cellular vitality.

**Figure 3.**
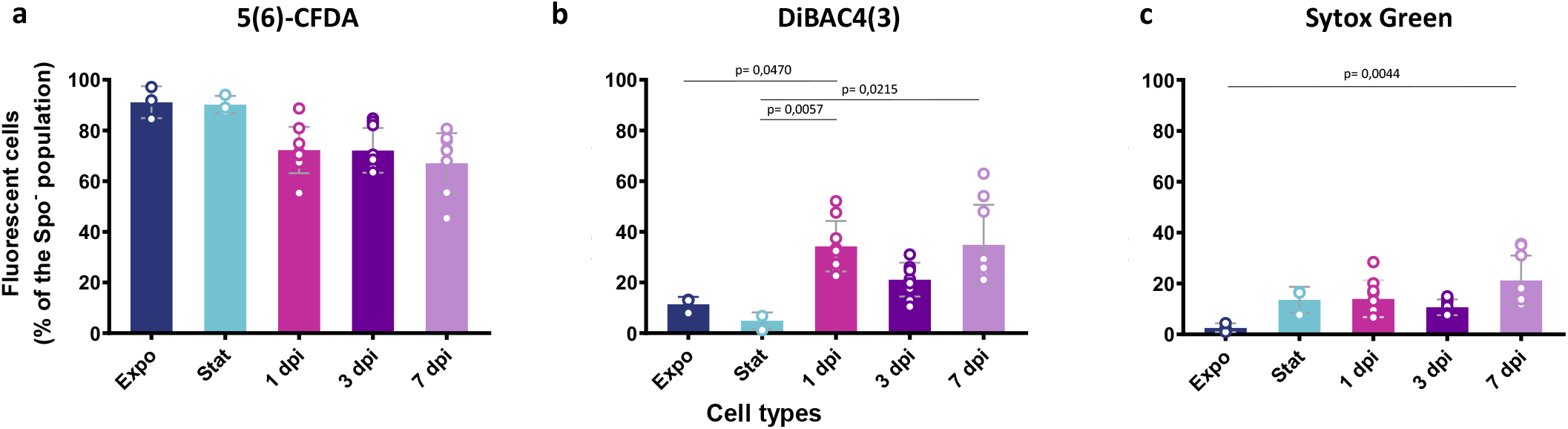
Assessment of the vitality status of the non-sporulated bacteria during infection. Flow cytometry analysis of Bt (pP*spoIIQ’mCherry*) cells grown in LB medium and harvested in exponential (OD_600_ = 1, dark blue) or stationnary phase (OD_600_ = 8, light blue), or extracted from *G. mellonella* cadavers at 1 day (pink), 3 (dark purple) and 7 days (light purple) post-infection. The samples were incubated for 10 min in the dark at RT with 5 µM 5(6)-CFDA **(a)**, 0,5 µM DiBAC4(3) **(b)** and 0.5 µM Sytox Green **(c)** before being subjected to analysis. The percentage of marked cells among non-sporulating bacteria discriminated in cytograms (as described in the Materials and Methods section) is presented as a function of time. Unmarked and heat-shocked bacteria were used as controls. Each symbol represents the data relative to bacteria extracted from one larva. The data are the result of three independent experiments and the error bars show the standard deviation from the mean. Statistically significant differences are indicated by black bars with the p-value obtained after a Kruskal-Wallis Test followed by a Dunn’s multiple comparison test.

### Protein production induction is delayed in non-sporulated late-infection stage bacteria

To further assess the physiological state of the 7 dpi bacteria, we examined the metabolic activity of these cells by recording their protein synthesis ability. We designed a strain harboring the transcriptional fusion P*x’gfp_Bte_* between a xylose-inducible promoter and a Gfp-encoding reporter gene associated to the sporulation reporter P*spoIIQ’mCherry*. We assessed the ability of the Spo^-^ cells to synthesize proteins by incubating them in conditioned HCT medium (HCTc) supplemented with xylose for 1 and 2 h and measuring the proportion of GFP-positive and mCherry-negative bacteria using flow cytometry (fig 4) coupled to microscopy observations (fig S5). Expo cells and Stat cells harvested from HCT cultures at 30°C, as well as bacteria extracted from insect cadavers at different time points (1, 3 and 7 dpi) were subjected to this treatement. The results show that at the start of induction, almost no bacteria expressed the P*x’gfp_Bte_* fusion in any of the samples, except for about 6% of the Stat and 1 dpi cells (fig 4). However, approximately 70% of the Spo^-^ Expo, Stat, 1 and 3 dpi cells, synthesized Gfp at 1 h post-induction. At 2 h post-induction the proportion of bacteria expressing *gfp_Bte_* in these samples slightly increased to 80%. Interestingly, the 7 dpi bacteria presented a different induction profile. At 1 h post-induction, less than 15% of the Spo^-^ bacteria were fluorescent, whereas at 2 h post-induction 15% to 75% of these cells were able to produce Gfp. This result indicates that the 7 dpi bacteria retain their ability to synthesize proteins, albeit with a greater time delay and variability between samples than for the other bacterial conditions assayed. This indicates that non-sporulating bacteria undergo physiological changes during the late infection stages suggestive of a metabolic slow-down.

**Figure 4.**
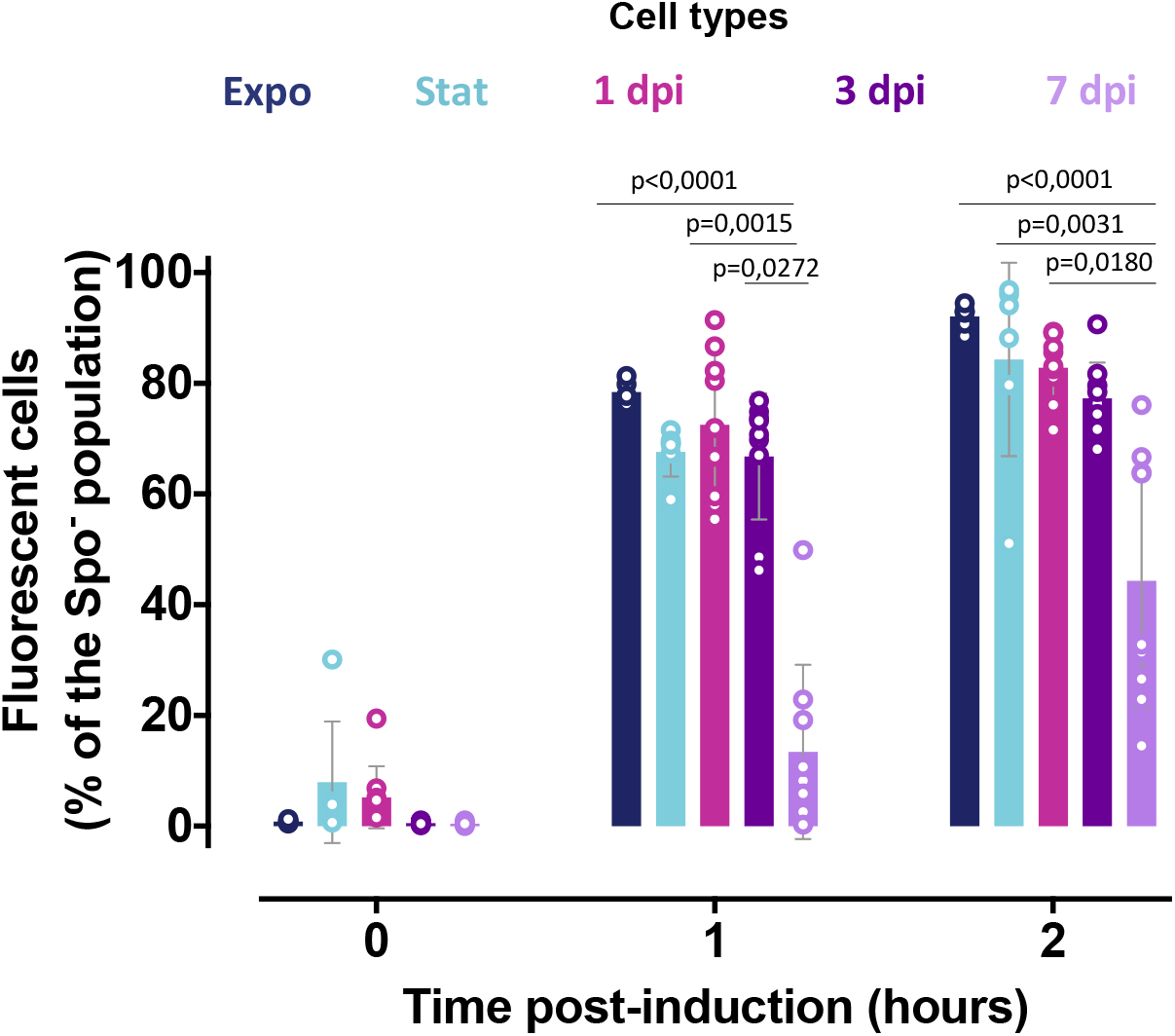
Induction of *gfp* expression in non-sporulated bacteria during infection. Flow cytometry analysis of Bt (pP*x’gfp_Bte_-*P*spoIIQ’mCherry*) cells grown in HCT medium and harvested in exponential (OD_600_ = 1, dark blue) or stationnary phase (OD_600_ = 6, light blue), or extracted from *G. mellonella* cadavers at 1 day (pink), 3 (dark purple) and 7 days (light purple) post-infection. The bacteria were resuspended in conditioned HCT medium supplemented with 25 mM xylose. Aliquots were analyzed at the time of inoculation and 1 and 2 h after the start of the induction. Green-fluorescent cells among the non-sporulating bacteria were discriminated in cytograms as described in the Materials and Methods section. Each symbol represents the data relative to bacteria extracted from one larva. The data are the result of three independent experiments and the error bars show the standard deviation from the mean. Statistically significant differences are indicated by black bars with the p-value obtained after a Kruskal-Wallis Test followed by a Dunn’s multiple comparison test.

### Growth recovery of late-infection stage bacteria is impaired on rich medium, in contrast to insect medium

We performed a colony-size assay to determine whether bacteria extracted at late time points from insect cadavers were able to replicate when exposed to fresh growth medium. Expo cells, Stat cells, spores harvested from 48 h-LB cultures at 30°C and bacteria extracted from insect cadavers at 1, 3 and 7 dpi were plated onto LB agar. Plates were photographed after 16 hours of incubation at room temperature (fig 5.a) and colony size was determined as described in the Materials and Methods section. The results show that colonies resulting from 3 and 7 dpi bacteria, as well as *in vitro*-prepared spores, were smaller than those resulting from Expo, Stat and 1 dpi cells (fig 5.a). This indicates that late-infection stage bacteria recover slower than bacteria sampled at an earlier infection stage or than *in vitro*-grown bacteria. This recovery is similar to that of *in vitro*-generated spores. The results might reflect a delay of growth due to the germination of the spores present at 3 dpi or after. However, all the colonies were smaller and therefore include the Spo^-^ cells, unless they are unable to form a colony. After longer incubation times, all the samples displayed similar colony-size (data not shown). In order to determine more precisely the behavior of the bacterial subpopulations, we also examined the growth of 7 dpi bacteria at the single cell level using time-lapse microscopy. Expo, Stat and 7 dpi bacteria were inoculated on LB agarose strips at 30°C and the behavior of Spo^-^ cells was followed during 5 hours (fig 5.c and 5.d). About 8 to 16% of the inoculated bacteria lysed regardless of the sample. Approximately 80% of Expo and Stat cells were able to divide whereas the majority of 7 dpi cells were inactive (fig 5.b). We observed elongation and growth for less than 20% of the 7 dpi bacteria (fig 5.c). These results show that 7 dpi bacteria rarely divided over a 5 h-time-lapse experiment which is in agreement with the slower growth recovery phenotype observed on LB plates.

**Figure 5.**
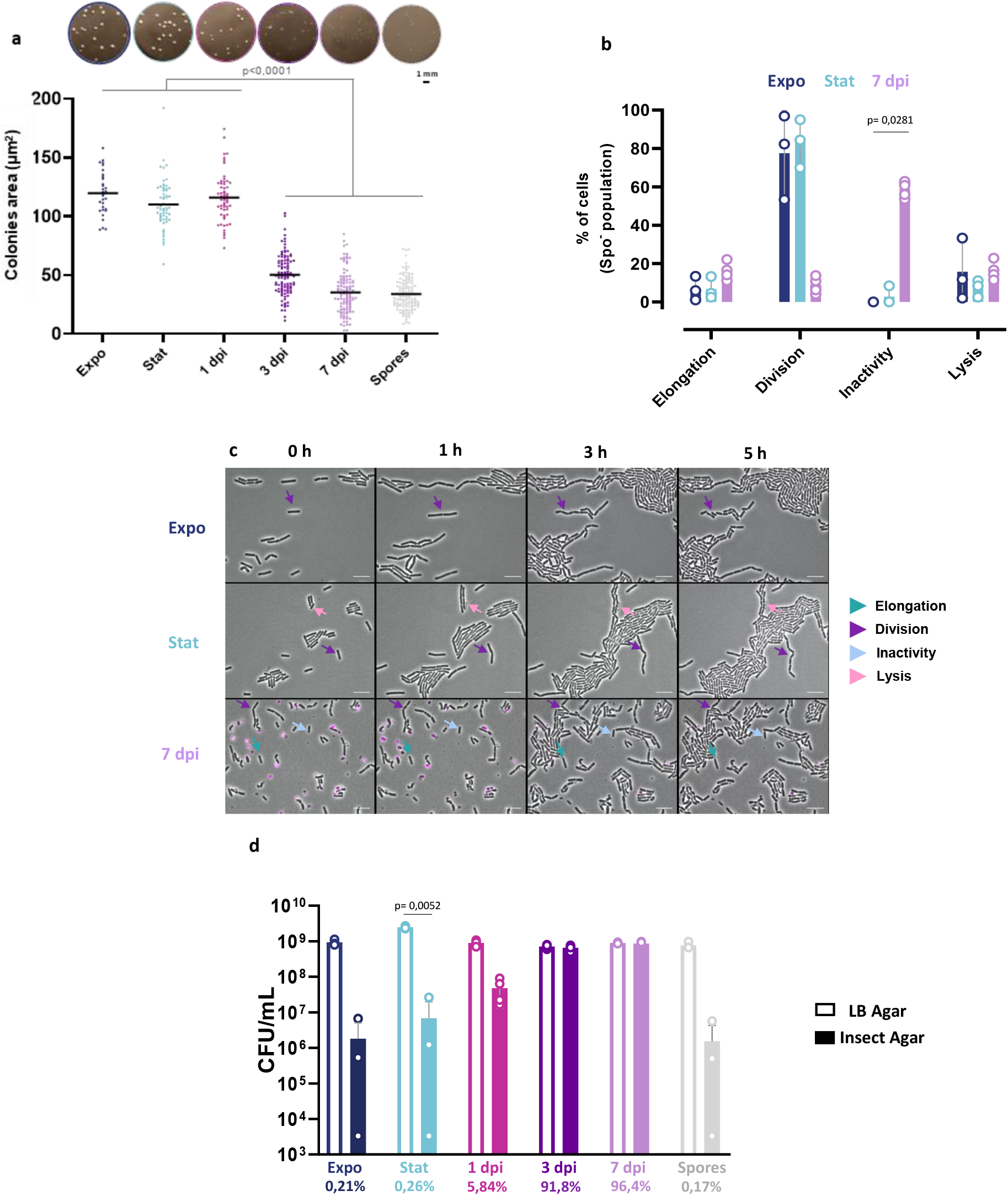
Recovery assessment of the bacteria extracted from insect cadavers. **a.** Colony size analysis of Bt (pP*spoIIQ’mCherry*) cells grown in LB medium and harvested in exponential (OD_600_ = 1, dark blue) or stationnary phase (OD_600_ = 8, light blue), or extracted from *G. mellonella* cadavers at 1 day (pink), 3 (dark purple) and 7 days (light purple) post-infection, or of an *in vitro* spore preparation in LB medium (grey). The upper panels show colonies photographed under a binocular stereo microscope (see details in the Materials and Methods section), 16 hours post-plating on LB agar. The scale bar represents 1 mm. Colony size was then measured using ImageJ as described in the Materials and Methods section. At least 145 cells were measured for each condition. Each symbol on the lower graph represents one colony. The data are the results of three independent experiments and the black horizontal bars indicate the mean. Statistically significant differences are indicated by black bars with the p-value obtained after a Kruskal-Wallis Test followed by a Dunn’s multiple comparison test. **b.** Time-lapse microscopy analysis performed on bacteria spotted onto LB agarose pads. One picture was taken per hour, during 5 hours. Each non-sporulating cell was affected to each of the listed categories and the percentage of cells for each category was calculated. The data are the result of three independent experiments. At least 145 cells were counted and the error bars show the standard deviation from the mean. Statistically significant differences are indicated by black bars with the p-value obtained after a Kruskal-Wallis Test followed by a Dunn’s multiple comparison test. **c.** Time-lapse microscopy pictures with the time scale represented above each picture. Bacteria were false colored in pink for Spo^+^cells. Arrows point to representative bacteria of the categories listed in b. The scale bar represents 10 μm. These results are representative of three independent experiments. **d.** Growth recovery on Insect agar medium. The bacteria were spotted onto Insect agar (filled bars) or LB agar plates (open bars). The bacteria were incubated at RT and CFU were numerated the next day of plating on the latter and 3 days later on the former. Each symbol represents bacteria extracted from one larva or originating from one culture. The percentage of CFU counted on LB versus Insect agar is indicated below each cell type. The data are the result of four independent experiments and the error bars show the standard deviation from the mean.

In order to determine if the late-infection stage bacteria were better adapted to a medium closer to the insect cadaver environement, we performed a recovery assay on an Insect agar medium prepared as described in the Materials and Methods section. Expo cells, Stat cells, spores harvested from 48 h-LB cultures at 30°C and bacteria extracted from insect cadavers at 1, 3 and 7 dpi were plated onto Insect agar plates. Figure 5.d shows that there is a drastic drop in the numbers of CFU/mL resulting from the plating of Expo and Stat cells, as well as spores, on Insect agar compared to LB. Less than 0.3% of these bacteria survived on this medium. About 6% of the 1 dpi population was able to grow on this medium compared to LB. In sharp contrast, the number of CFU/mL for 3 and 7 dpi cells was similar on both media. Taken together, these results suggest that the insect cadaver is a hostile environment for bacteria and that adaptation is required to survive in these conditions.

### Transcriptomic profile of late-infection B. thuringiensis

To determine the expression profile of the Spo^-^/Nec^-^ cells and begin to understand how these bacteria persist in the cadavers, we performed a global *in vivo* transcriptional analysis at a late stage of infection using an RNA-Sequencing approach. We chose to extract RNA from the total population at the 7 dpi time-point that presented very few Nec^+^ bacteria (fig 2) and we verified that RNA was only poorly extracted from spores, thus preventing a biais in our analysis (see Materials and Methods section). We compared the transcriptome of 7 dpi bacteria to Expo and Stat bacteria grown in LB medium. The transcriptome of the Stat and Expo bacteria grown in LB medium was also compared to identify genes differentially expressed in the insect specifically. As presented in fig 6.a, the principal component analysis shows that the 7 dpi bacteria differed substantially from both Expo and Stat cells, indicating that these cells are in a different state. The results show that 484 and 203 genes were specifically up-regulated and down-regulated, respectively, in the 7 dpi cells when compared to both Expo and Stat bacteria (fig 6.b). We found that most of the genes belonging to the NprR regulon were down-regulated when compared to Stat cells but up-regulated when compared to Expo cells (Table S4). This is in accordance with the fact that a small proportion of bacteria are Nec^+^ at 7 dpi (fig 1) while necrotrophism is repressed in exponential phase and activated during stationary phase *in vitro* [28]. The data also show that most sporulation and germination genes were not differentially expressed, indicating that these two processes were not triggered in the subpopulation of interest (Table S5, S6).

**Figure 6.**
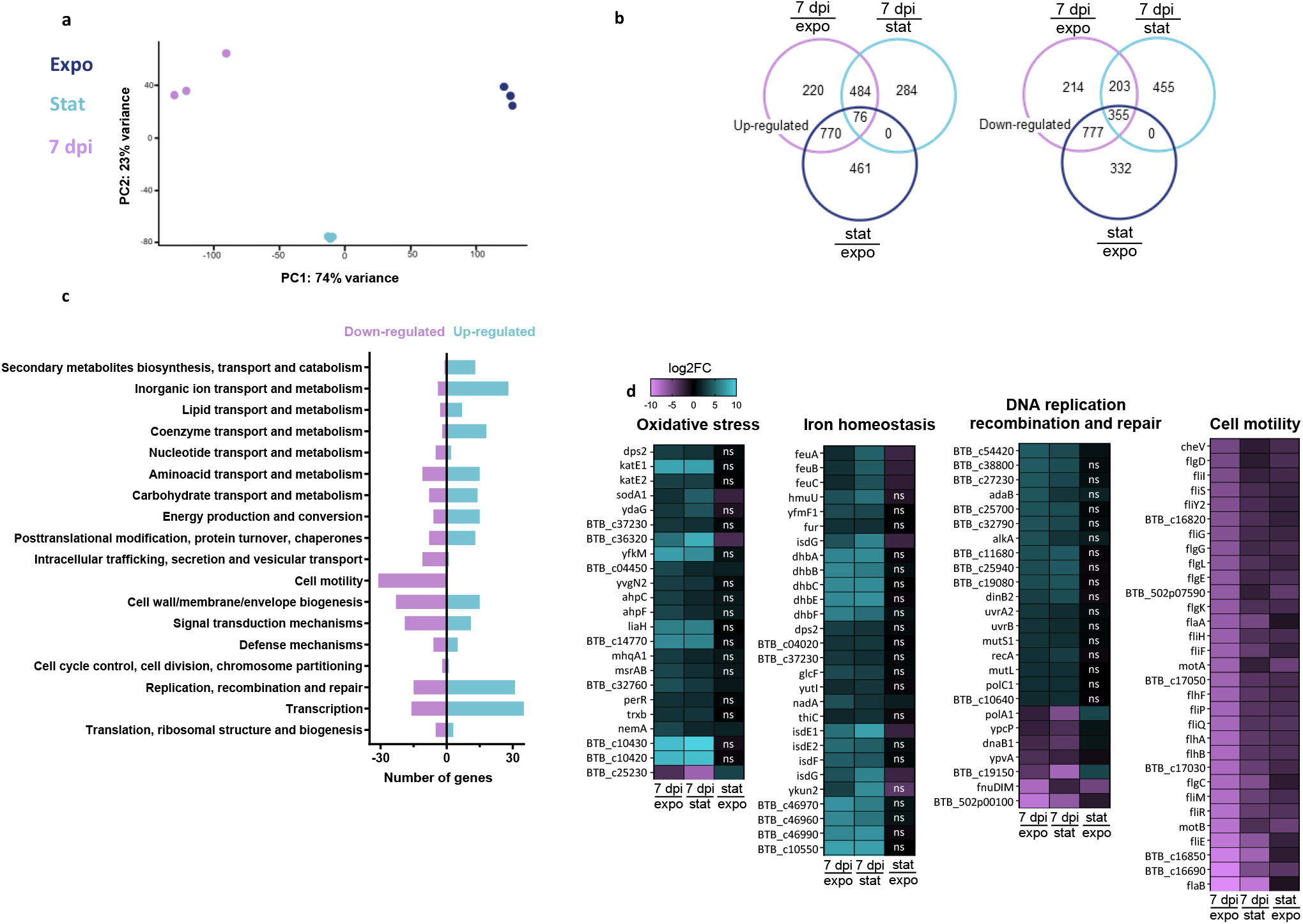
Transcriptomic analysis of bacterial cells extracted from insect cadavers at 7 days post-infection. **a.** Principal component analysis plot of the different samples: exponential growth phase cells (dark blue), stationary growth phase cells (light blue), cells extracted from insect cadavers at 7 dpi (purple). Samples were plotted against the first two principal components calculated from the gene expression values. The axes labels indicate percentage of total variance that is explained by each component. **b.** Venn diagrams indicating the number of up-regulated and down-regulated genes in cells extracted from insect cadavers at 7 dpi compared to exponential growth phase cells (purple) or stationary growth phase cells (light blue) and in stationary growth phase cells compared to exponential growth phase cells (dark blue). The genes taken into account present a log2FC≤-2 or log2FC≥2 and an adjusted p-value≤0,01. **c.** COG analysis of genes expressed in cells extracted from insect cadavers at 7 dpi compared to exponential growth phase cells and stationary growth phase cells. Blue bars indicate up-regulated genes with a log2FC≥2, purple bars indicate down-regulated genes with a log2FC≤-2. The considered genes have an adjusted p-value≤0,01. Genes with unknown functions are not represented. **d.** Heatmaps of genes expressed in cells extracted from insect cadavers at 7 dpi compared to exponential growth phase cells (left column), stationary growth phase cells (mid column) or in bacteria harvested in stationary growth phase compared to exponential growth phase (right column). Selected categories of differentially expressed genes are shown: oxidative stress, iron homeostasis, DNA replication recombination and repair as well as cell motility. Purple indicates low expression (log2FC ≤-2) and blue indicates high expression (log2FC≥2). ns indicates an adjusted p-value>0,01.

Most of the up-regulated genes in the 7 dpi cells are involved in inorganic ion transport and metabolism, transcription, replication, recombination and repair, whereas the down-regulated genes are mostly involved in cell motility, signal transduction mechanisms and cell wall/membrane/envelope biogenesis (fig 6.c), the latter indicating a decrease in the growth process. More than 300 differentially expressed genes were assigned to an unkown function. Differentially expressed genes with the highest or lowest log2FC values were found in the oxidative stress, iron homeostasis, DNA replication, recombination and repair and cell motility categories (fig 6.d). Interestingly, all motility genes were down, all iron homeostasis genes were up, and all oxidative stress genes were up (except for one), indicating that these mechanisms are key for late infection stage survival.

### Oxidative stress genes are specifically expressed during late-infection, in contrast to iron homeostasis genes expressed at all times

To examine the expression profile of selected genes identified in the RNA-Seq analysis and determine which of these were specific to late infection stage survival, we constructed strains harboring a transcriptional fusion between the promoter region of the gene of interest and the *gfp_Bte_AAV* reporter gene, associated to the sporulation reporter P*spoIIQ-mCherry*. We chose genes with the highest log2FC values obtained by the differential gene expression analysis in the iron homeostasis and oxidative stress resistance categories. We followed the expression profile of representative genes from these categories such as *ykun2* (a flavodoxin that replaces ferredoxin under conditions of iron limitation), *isdE1* (involved in heme scavenging), *dhbA* (the first gene of the *dhb* operon that specifies the biosynthesis of the siderophore bacillibactin involved in the chelation of ferric iron from the surrounding environment), *BTB_c10430* (annotated as *sigX* by KEGG orthology), *katE1* (a catalase and general stress protein) and *sodA1* (a superoxide dismutase and general stress protein) [26]. P*ykun2’gfp_Bte_AAV* and P*isdE1’gfp_Bte_AAV* were expressed from the start of the infection, and the percentage of fluorescent cells among the Spo^-^ cells was around 30% (fig 7.a and 7.b). Similarly, P*dhbA’gfp_Bte_AAV* was expressed in approximately 40% of the cells among the Spo^-^ bacteria in the 1, 3 and 7 dpi samples (fig 7.c). For the reporter P*BTB_c10430’gfp_Bte_AAV* less than 20% of the 1 and 3 dpi Spo^-^ cells were Gfp-positive. At 7 dpi, around 40% of the Spo^-^ cells produced Gfp, indicating a specific activation of *BTB_c10430* during in late infection survival (fig 7.d). P*katE1’gfp_Bte_AAV* and P*sodA1’gfp_Bte_AAV* were specifically expressed in 7 dpi cells with about 65% of green fluorescent Spo^-^ cells for the former and 80% for the latter (fig 7.c). As expected, none of these reporters were expressed in Expo and Stat cells. Altogether, these results indicate that iron homeostasis is important from the start of the infection whereas an oxidative stress response is mounted as the bacteria spend more time in the cadaver environment.

**Figure 7.**
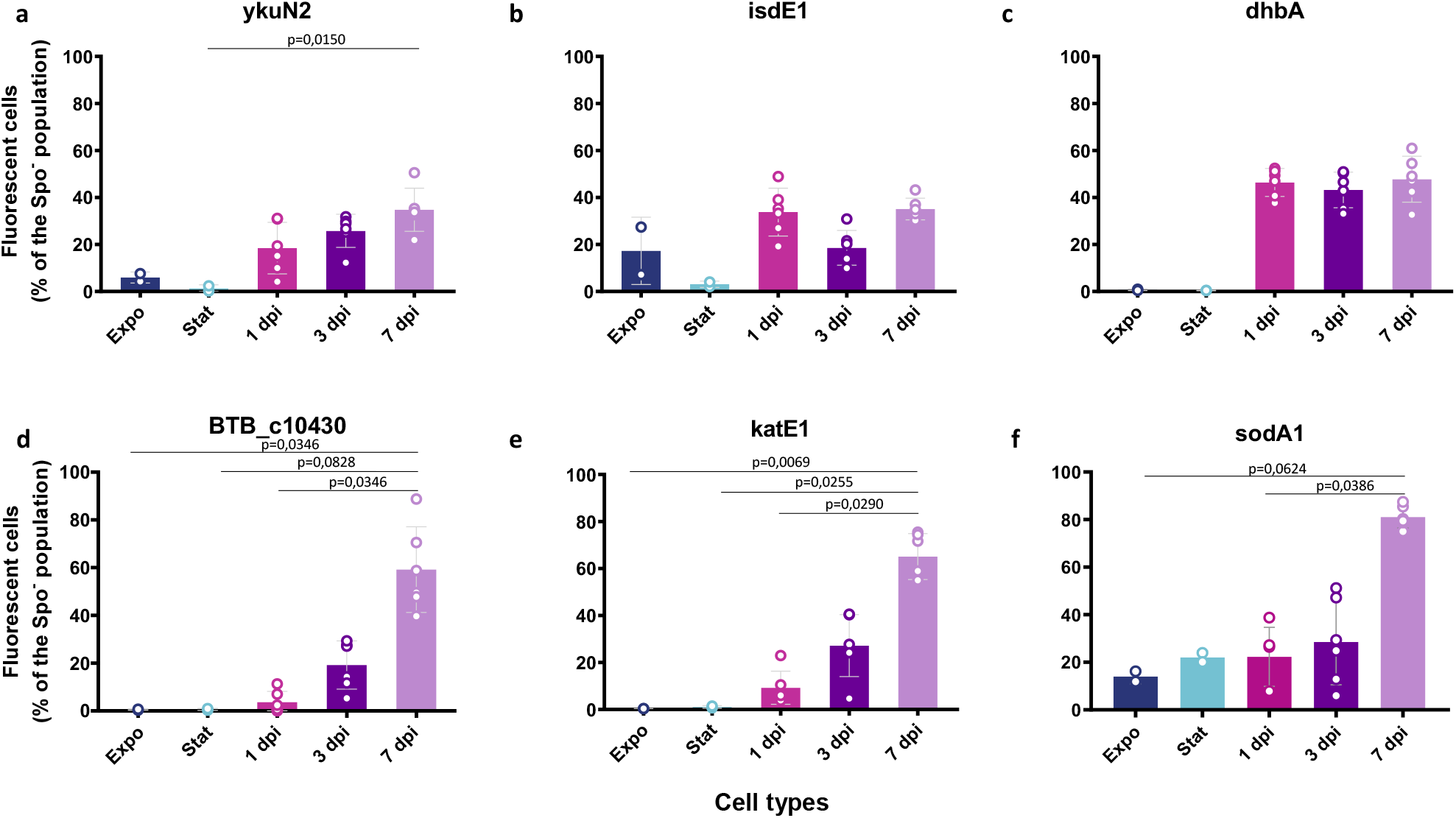
Iron homeostasis and oxidative stress resistance promoter activities during infection. Flow cytometry analysis of Bt (pP*ykuN2’gfp_Bte_AAV-*P*spoIIQ’mCherry*) (**a**), Bt (pP*isdE1’gfp_Bte_AAV-*P*spoIIQ’mCherry*) (**b**), Bt (pP*dhbA’gfp_Bte_AAV-*P*spoIIQ’mCherry*) (**c**), Bt (pP*BTB_c10430’gfp_Bte_AAV-*P*spoIIQ’mCherry*) (**d**), Bt (pP*katE1’gfp_Bte_AAV-*P*spoIIQ’mCherry*) (**e**) and Bt (pP*sodA1’gfp_Bte_AAV-*P*spoIIQ’mCherry*) (**f**) cells grown in LB medium and harvested in exponential (OD_600_ = 1, dark blue) or stationnary phase (OD_600_ = 8, light blue), or extracted from *G. mellonella* cadavers at 1 day (pink), 3 (dark purple) and 7 days (light purple) pi. Green fluorescent cells among the non-sporulating bacteria were discriminated in cytograms as described in the Materials and Methods section. Each symbol represents the data relative to bacteria extracted from one larva. The data are the result of two independent experiments and the error bars show the standard deviation from the mean. Statistically significant differences are indicated by black bars with the p-value obtained after a Kruskal-Wallis Test followed by a Dunn’s multiple comparison test.

## DISCUSSION

Our study reports the characterization of a physiological state that procures the sporulating entomopathogenic Gram-positive bacterium *B. thuringiensis* specific properties allowing its persistence in the host environment in a non-sporulated form. We analyzed the phenotypic properties and transcription profile of the Spo^-^ *B. thuringiensis* cells in the context of an infection in *G. mellonella* larvae. These larvae are natural hosts for the *B. thuringiensis* species, but they are also widely used as an alternative to mammalian models of infection to study bacterial virulence and host colonization [30, 31].

The RNA-Seq analysis showed that the motility genes were down-regulated in 7 dpi bacteria compared to *in-vitro*-grown cells, and that the motility repressor-encoding gene *mogR* [32] was up-regulated in 7 dpi bacteria compared to exponentially growing cells. Similarly, the biofilm-associated genes *calY*, *tasA* and the *eps2* locus [33, 34], were up-regulated compared to exponentially growing cells. These data suggest that the late-infection stage bacteria form a biofilm which might confer them an advantage for long-term survival considering the resistance properties of these structures [35]. The most represented categories among the up-regulated transcripts in 7 dpi bacteria compared to *in vitro*-grown cells are the oxidative stress response, DNA recombination, replication and repair in addition to iron homeostasis. The expression of uptake systems for iron, as well as biosynthesis and uptake of the endogenous siderophore bacillibactin was shown to be activated by iron limitation in *B. cereus* [36]. During an infection, the host is considered to be an iron-depleted environment and bacteria need to deploy strategies to scavenge this element, such as the activation of the Fur regulon [37, 38]. This is consistent with the activation of the iron homeostasis genes *dhbA* and *isdE1*, belonging to this regulon, during all stages of the infection. Iron is essential for the function of various proteins and iron homeostasis was previously reported as being important for *B. cereus* pathogenesis in the first stages of *G. mellonella* infection by oral gavage [39]. However, this metal is also a potential hazard in combination with compounds such as H_2_O_2_ because the reaction between the two products generates reactive oxygen species (ROS) that can be harmful by damaging cellular components such as DNA or proteins with [Fe-S] clusters [40]. Several studies have linked iron homeostasis and DNA repair to a response to oxidative stress and showed the upregulation of these gene categories upon H_2_O_2_ exposure [41, 42], and it was recently reported that H_2_O_2_ was produced in the hemolymph of *G. mellonella* during *Salmonella enterica* infection [43]. Moreover, the melanization defense reaction that occurs after the insect infection liberates ROS and might last for several days [44–46]. Aerobic metabolism also generates endogenous oxidative stress if O2 is not completely reduced during respiration and a secondary oxydative stress response can also be triggered when *Bacillus* encounter unfavorable conditions in an aerobic environment [47]. All these stresses are counteracted by the action of proteins such as catalases (Kat), superoxide dismutases (SodA) or DNA-binding ferroxidases (Dps) [48, 49], all of which are overexpressed in 7 dpi bacteria. RNA-Seq analysis also revealed that the BTB_c10430 locus, annotated as encoding the SigX protein [26], is specifically up-regulated in 7 dpi bacteria. SigX belongs to the extracytoplasmic function (ECF) sigma factor family that helps maintain cell envelope homeostasis and activate resistance to agents that can compromise the integrity of the envelope, the first defense against environmental threats [50]. A *B. subtilis sigX* deletion mutant was shown to be sensitive to oxidative stress [51]. Further investigations should examine the involvement of this regulatory protein in the maintenance of *B. thuringiensis* in insect cadavers. It would also be informative to determine if the oxidative stress response triggered in the late-infection stage bacteria is due to the cadaver environment, to an excessive iron uptake or to a combination of the two and to understand the role of the oxidative stress response in the resistance of *B. thuringiensis* during long-term infection.

*B. thuringiensis* is able to overcome the immune defenses of *G. mellonella*, to multiply and to completely invade its host in less than 24 hpi [11,52,53], (this study and unpublished). It was shown that the bacteria engage in necrotrophism after the death of the larva by activating the NprR regulon that includes genes encoding chitinases, proteases and oligopeptide permeases [11, 15]. Activation of this regulon seemed to last throughout the duration of the infection [15] and an *nprR* deletion mutant was unable to survive in the cadaver suggesting that the regulon was required for long-term survival. However, we showed here that necrotrophism is a transient state limited to the first stages of survival in the cadaver. We propose that this state allows *B. thuringiensis* to scavenge the nutrients present in the cadaver shortly after the insect death, then the bacteria enter different pathways, potentially triggered by a stress signal such as nutrient depletion, that allow for their persistence in the host. A subpopulation enters the differentiation pathway leading to sporulation, while another engages in a metabolic slow-down that accentuates with the increase of exposure time to the cadaver environment. This subpopulation also showed resistance to the cadaver environment which proved hostile for *in vitro*-grown bacteria as well as *in-vitro* generated spores.

Altogether, these results show that non-sporulated late-infection stage *B. thuringiensis* are resilient cells able to resist in a harsh environment and suggest that this adaptation relies mainly on the activation of stress resistance genes as well as entering a metabolic slow-down. The fitness advantages conferred by the phenotypic heterogeneity of the *B. thuringiensis* population during infection and, in particular, by the co-existence of spores and of the non-sporulated form able to survive for a prolonged period in a host cadaver, still need to be elucidated. Studying the mechanisms that govern this state will provide valuable fundamental knowledge about the lifecycle of these bacteria and might lead to the development of new strategies to combat sporulating pathogenic species.

## Supporting information

TOUKABRI et al_Supplemental

TOUKABRI et al_TableS4

TOUKABRI et al_TableS5

TOUKABRI et al_TableS6

## ACKNOWLEDGEMENTS

The authors are indebted to Christophe Buisson for insect rearing. This work has benefited from the facilities and expertise of the high throughput sequencing core facility of I2BC (Centre de Recherche de Gif – http://www.i2bc.paris-saclay.fr/) and the authors thank Delphine Naquin for primary analysis. We are grateful to Tarek Toukabri, Agnès Réjasse and Sébastien Gélis-Jeanvoine and Yan Jaszczyszyn for their help with the RNA-Seq experiment and analysis. The CyFlow Space flow cytometer was funded by the DIM Astrea (French regional program Ast11 0137). HT was the recipient of a doctoral grant from the ministry of French higher education, delivered by the Doctoral School ABIES at AgroParisTech–Université Paris Saclay. This work was supported by the MICA department of INRAE (French National Research Institute for Agriculture, Food and Environment).

