## Supplementary material for "A sporulation-independent way of life for *Bacillus thuringiensis* in the late stages of an infection": TOUKABRI et al_Supplemental

Hasna Toukabri, Didier Lereclus and Leyla Slamti*

Micalis Institute, INRAE, AgroParisTech, Université Paris-Saclay, 78350 Jouy-en-Josas, France

The authors declare that there are no competing interests in relation to the work described.

**MATERIALS AND METHODS**

**DNA manipulations**

Plasmid DNA was extracted from *E. coli* by a standard alkaline lysis procedure, using a Promega kit (Promega,Madison, Wisconsin USA). Restriction enzymes, T4 DNA ligase, Standard Taq DNA polymerase and Phusion high-fidelity DNA polymerase were purchased from New England Biolabs (Ipswich, MA, USA) and used as recommended by the manufacturer. The oligonucleotide primers (Table S3) used for PCR amplification were synthesized by Eurofins Genomics (Nantes, France). PCR was performed with a 2720 Thermak cycler (Applied Biosystems). All constructs were systematically verified by PCR followed by sequencing of the region of interest. Nucleotide sequences were determined by Eurofins Genomics (Köln, Germany).

**Flow cytometric analysis**

For GFP-, 5(6)-CFDA-, DiBAC4(3)- and Sytox Green-based fluorescence, a solid blue-laser emitting at 488 nm was used, combined to a 500-nm long pass dichroic mirror and a 527-nm band pass filter (512–542) (FL1 Channel). For mCherry-based fluorescence, a solid yellow-laser emitting at 561 nm was used, combined to a 610-nm long-pass filter (FL4 channel). The analyses were performed using logarithmic gains and detector settings, adjusted on a sample of reporterless cells, to define cellular autofluorescence. Gating on FSC⁄SSC was used to discriminate bacteria from the background. For each sample, 20000 gated events were measured. Data were collected with the FlowMax software (Sysmex Partec, France) and analyzed with the Weasel 3.3.3 software (WEHI, USA).

To identify positive and negative populations on FL1/FL4 bi-parametric cytograms, we applied the 98% division line for each fluorescent marker, i.e. we set the threshold on the reporterless strain so that 98% of the population gave a fluorescence intensity below the threshold. Bacteria with a fluorescence signal above the threshold were considered positive. We cannot exclude that for a reporter expressed at a low level, a few positive cells might have a fluorescent intensity similar to that of the reporterless cells and be included in the negative population.

**RNA-Extraction**

2 mL aliquots were sampled from LB cultures at OD_600_=1 (exponential phase growth) and OD_600_=8 (stationary phase growth) and immediately mixed with an equal volume of RNA-later (Invitrogen, Eugene, OR, U.S.A.). Bacteria from insect cadavers were crushed in 1 mL of RNA-later and vortexed. All samples were stored at 4°C overnight to allow thorough penetration of RNA-later.

For bacteria extracted from insect cadavers, the liquid fraction was transferred to a new tube and centrifuged for 10 min at 13000 rpm at 4°C. The fat pellicle and supernatant were discarded and the pellet resuspended in 750 µL of saline. The suspension was then filtered on a cotton pad in a 1 mL syringe to remove larvae debris. Then, all samples (from *in vitro* cultures and from insect cadavers) were centrifuged for 3 minutes at 13000 rpm at 4°C and the pellets were resuspended in 1 mL of Trizol (Invitrogen, Eugene, OR, USA). Resuspended bacteria were disrupted by adding silica beads (Biospec Products, Bartlesville, OK, USA) to the suspension and shaking in a Fastprep 24 (MP Biomedicals) for 45 s at 6.5 M/s twice. The supernatant was transferred to a clean tube and 100 µL of 1-bromo-3-chloropropane (Sigma-Aldrich, Saint-Louis, MO, USA) were added. The suspension was vortexed, incubated for 10 min at room temperature and centrifuged for 15 min at 13000 rpm at 4°C. The aqueous phase was then transferred to a clean tube to which 0.1 volume of sodium acetate and 0.7 volume of isopropanol were added. The suspension was mixed with 1 µL of glycogen (Thermo Fisher Scientific, Waltham, MA, USA) and incubated at -20°C for 1 h to precipitate nucleic acids. Nucleic acids were then pelleted by centrifugation (20 min at 13000 rpm at 4°C), washed twice with 75% ethanol, air-dried and resuspended in RNase-free water (Thermo Fisher Scientific, Waltham, MA, USA). Traces of contaminating DNA were removed by TURBO DNase treatment (Invitrogen, Eugene, OR, USA).

**RNA-Seq analysis**

To assign functional categories to the differentially expressed genes, we used a database constructed by Sébastien Gélis-Jeanvoine (unpublished). This database was constructed as follows: all the genes of *B. thuringiensis* 407 were functionally re-annotated using InterProScan [1], HHSearch [2] and EggNOG [3]. InterProScan v.5 was queried online through its SOAP API (http://www.ebi.ac.uk/Tools/webservices/services/pfa/iprscan5_soap) with the --goterms and --nopathways flags. GO terms were then parsed and used as three different annotations (process, function and component). HHSearch was used on our local cluster to query an HMM database prepared from the PDB70 database (2014-09-06 update), with a probability threshold of 95%. COG letters were assigned to our query genes by first aligning them with BLASTp (e-value threshold of 10^-2^) against the EggNOG v4.0 data base. The best hit for each query gene was then used to retrieve the cognate COG letter via an in-house script (EggnogGenome, <https://github.com/seb-ksl/EggnogGenome)>.

**FIGURES**

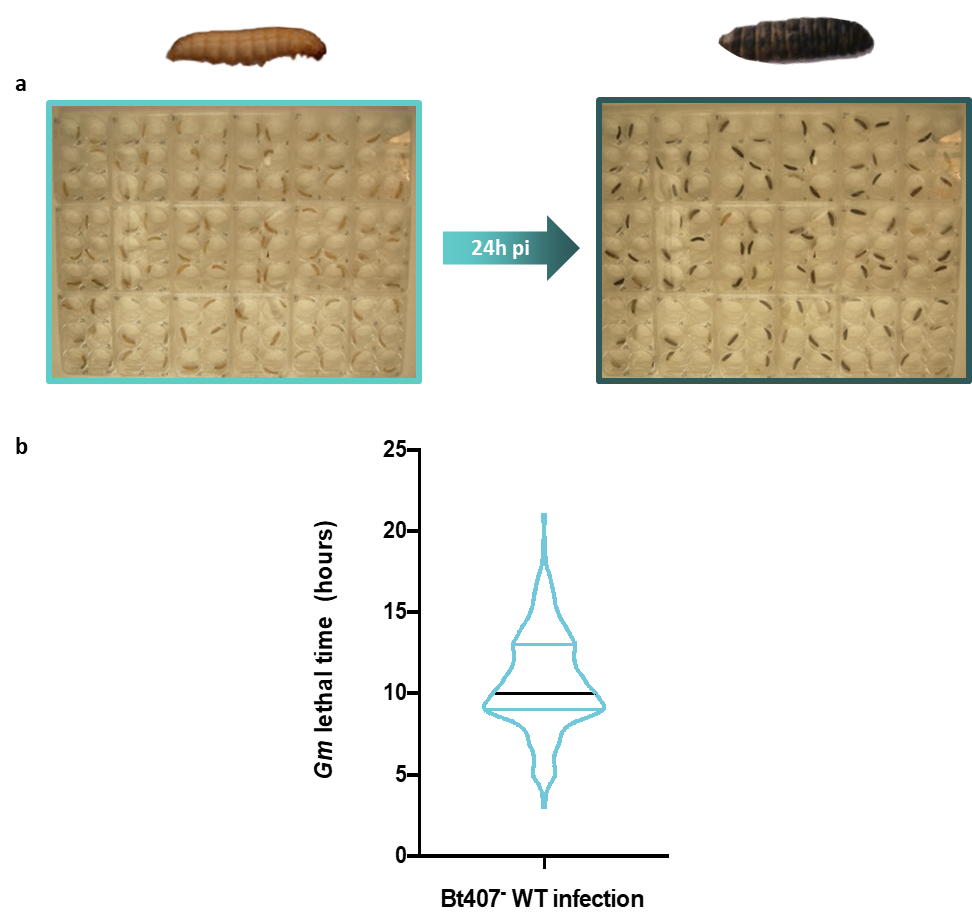

***G. mellonella* lethal time (hours)**

***B. thuringiensis* infection**

**24 hpi**

**Figure S1. Time-lapse photography set-up to determine *G. mellonella* lethal time.**

**a.** First picture shows infected larvae incubated in 6-well plates at 30°C under a Nikon CoolPix P1 camera on time-lapse photography mode to determine the time of death for each larva. Melanization as shown by the second picture at 24 hpi and absence of movement are required to consider a larva as dead. **b.** *G. mellonella* lethal time after infection with *B. thuringiensis* strain 407. A picture was taken every 10 min with the time-lapse photography set-up and lethal time post-infection is reported. Black line indicates the median, blue lines indicate the first and last quartile, n>150.

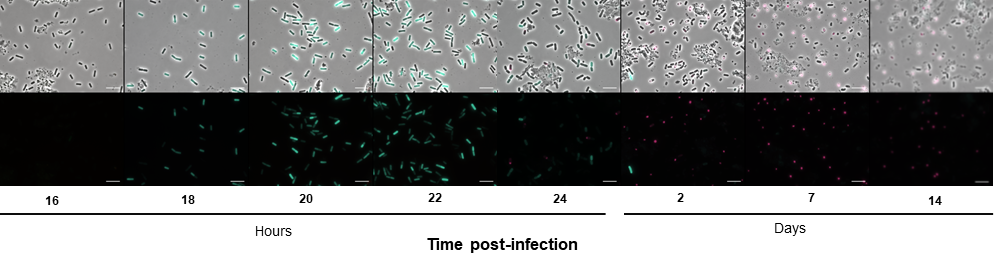

**Figure S2. Microscopy observations of the necrotrophism and sporulation promoters activity in bacterial cells during long-term infection.**

Bacteria were analyzed by fluorescence microscopy at the time points indicated. Upper panels merge between the phase contrast and epifluorescence images channels; lower panels, epifluorescence images. Cells were false colored in green for Nec^+^ cells and pink for Spo^+^ cells. The scale bars represents 10 μm.

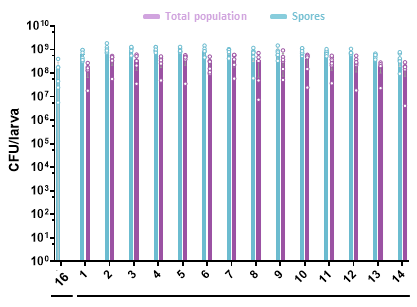

**Hours**	**Days**

**Time post-infection**

**Figure S3. Monitoring of *B. thuringiensis* survival during long-term infection.**

The total bacterial population (purple) and spores (blue) were numerated daily for 14 days after intrahemocoelic infection of *G. mellonella* with Bt (pP*nprA’gfp_Bte_AAV*-pP*spoIIQ’mCherry*). 6 larvae were crushed at the time points indicated and serial dilutions of the homogenate were directly plated onto LB agar for total population numeration. The bacterial suspensions were heated at 80°C during 12 minutes and plated onto LB agar to count heat-resistant spores. Each symbol represents bacteria extracted from one larva. The data are the result of two independent experiments and the error bars show the standard deviation from the mean.

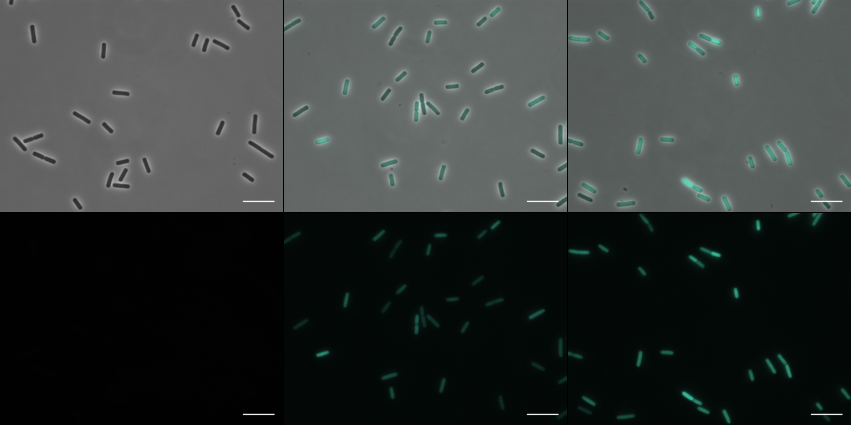

**Expo**

**Stat**

**1 dpi**

**3 dpi**

**7 dpi**

**0h**

**1h**

**2h**

**Time post-induction (hours)**

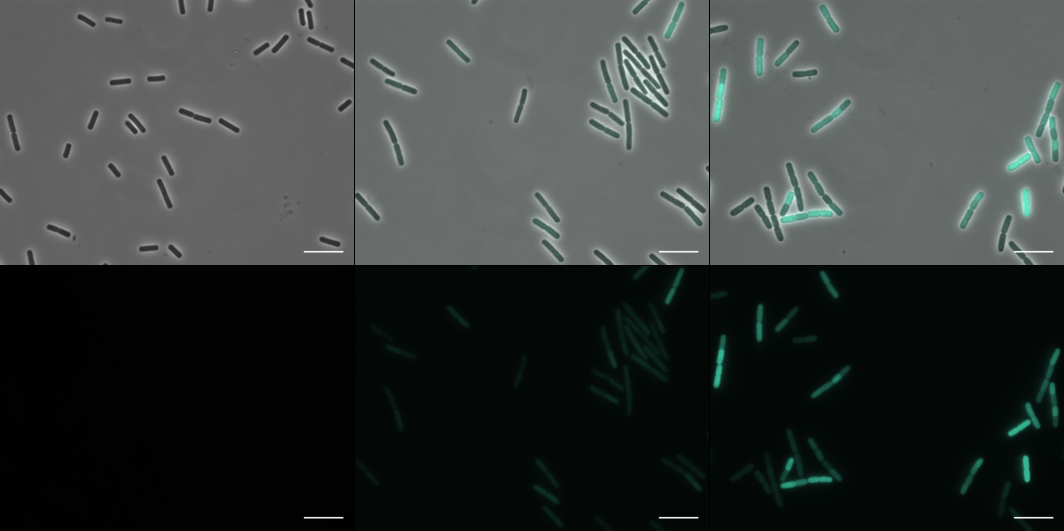

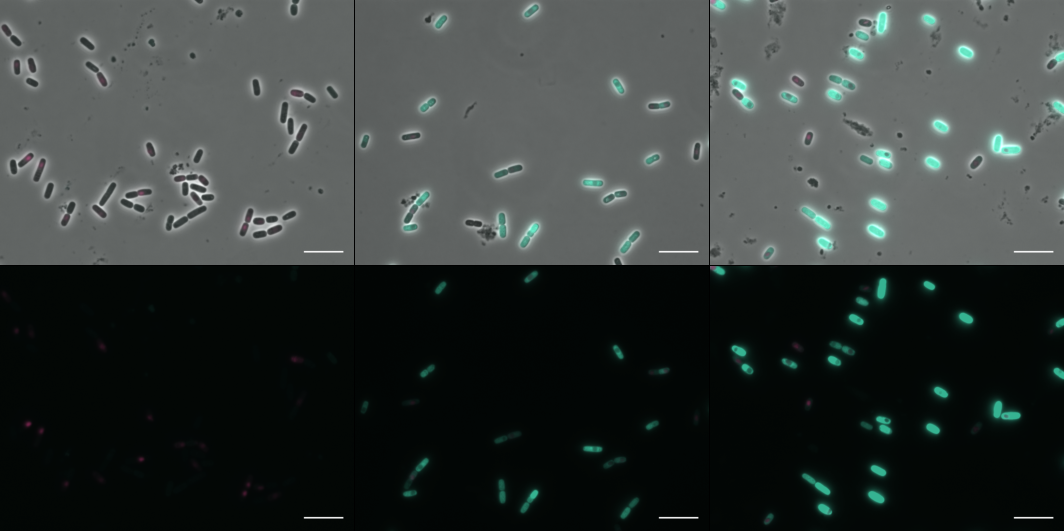

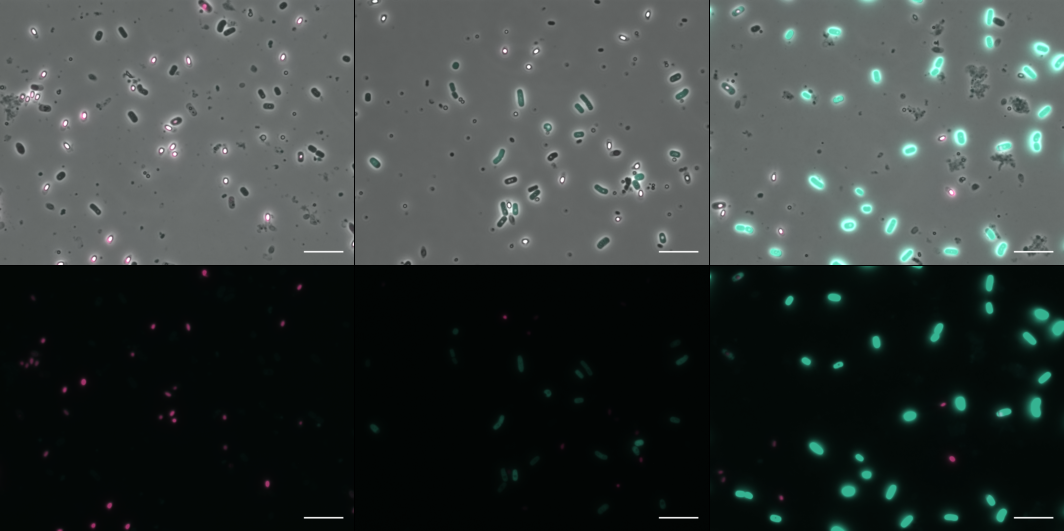

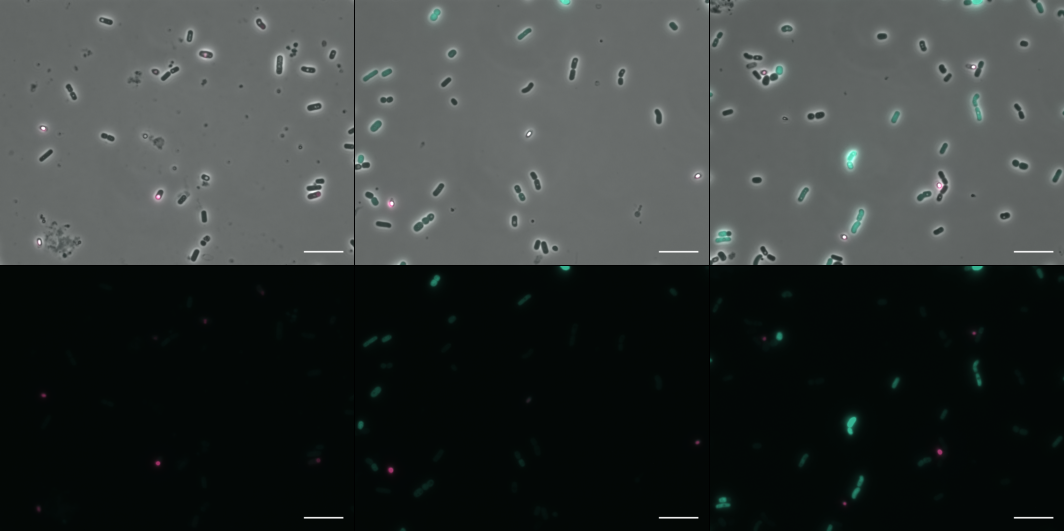

**Figure S6. Induction of *gfp* expression in non-sporulated bacteria**. Fluorescence microscopy images at the time after induction indicated above the pictures. Lower panels, epifluorescence images; upper panels, merge between the two channels. Cells were false colored in green for gfp-expressing cells and pink for Spo^+^ cells. The scale bar represents 10 μm. These results are representative of three independent experiments.

**TABLES**

**Table S1. Plasmids used in this study**

| **Name** | **Relevant information** | **Référence** |
| --- | --- | --- |
| pHT304 | Replicative multicopy *E. coli*/*B. thuringiensis* shuttle vector. | [4] |
| pHT304.18 | Replicative multicopy *E. coli*/*B. thuringiensis* shuttle vector. | [5] |
| pP*x’gfp_Bte_* | *B. thuringiensis* codon optimized *gfp* including 24 bp encoding the first eight amino acids of *comGA* [6], cloned in the pHT304’P*xyl+* vector with the modified RBS AGGAGG [7]. | [8] |
| pHT-*gfp_Bte_AAV* | Destabilized *gfp* using *ssrA* –AAV tag designated *gfpAAV* cloned into pHT304.18. | [8] |
| pP*nprA*’*gfpAAV* | The promoter region of the *nprA* gene was amplified by PCR from the chromosome of *B. thuringiensis* strain 407 using primer pairs P*nprA*-F-XbaI/P*nprA*-R-AscI and cloned between the XbaI and AscI restriction sites of pHT-*gfpAAV* to detect fluctuations in necrotrophism expression. | This study |
| pP*spoIIQ* | The promoter region of the *spoIIQ* gene was amplified by PCR from the chromosome of Bt 407 using primer pairs P*spoIIQ*-F-SphI/P*spoIIQ*-R-XbaI and cloned between the SphI and XbaI restriction sites of pHT304.18 to assess sporulation resulting into the construction p*PspoIIQ’*. This construction was prepared for future *mCherry* insertion and optimization by cloning the transcriptional terminator of phage lambda [9] using primer pairs Term-F-KpnI/Term-R-EcoRI between the KpnI and EcoRI restriction sites of p*PspoIIQ*’. We also added The STAB-SD [10] sequence for future optimization purpose. STABS-SD was amplified by PCR using primer pairs StabSD-F-XbaI/StabSD-R-BamHI and cloned between the XbaI and BamHI restriction sites of pP*spoIIQ*’TermLambda. We finally obtained pP*spoIIQ*’StabSD’TermLambda. | This study |
| pP*spoIIQ*’*mCherry* | *B. thuringiensis* codon optimized ComGAmCherry was amplified by PCR from Pp*x+*’*comGAmCherry* (unpublished) using primer pairs comGAmC-F-BamHI/mC-R-KpnI and cloned between the BamHI and KpnI restriction sites of pP*spoIIQ*’StabSD’TermLambda to create a stable transcriptional fusion to assess sporulation on a long-term scale. |  |
| pP*spoIIQ*’*mCherry*’-P*nprA*’*gfpAAV* | P*spoIIQ*’StabSD’*comGAmCherry*’TermLambda was amplified by PCR from pHT’P*spoIIQ*’StabSD’*comGAmCherry*’TermLambda using primer pairs P*spoIIQ*-F-SalI/Term-R-SphI and cloned between the SalI and SphI restriction sites of pP*nprA*’gfpAAV. This construction is used to monitor transient necrotrophism expression and sporulation during late stages of infection. | This study |
| pP*x+*’*gfp* | The transcriptional terminator of phage lambda was amplified by PCR using primer pairs Term-F-KpnI/Term-R-AscI-XbaI-EcoRI and cloned between the KpnI and EcoRI restriction sites of pP*x+*’*gfp* to obtain pP*x+*’*gfp*’*Term* | This study |
| pP*x+*’*gfp*’-P*spoIIQ*’*mCherry* | P*spoIIQ*’StabSD’*comGAmCherry*’TermLambda was amplified by PCR from pHT’P*spoIIQ*’StabSD’*comGAmCherry*’TermLambda using primer pairs P*spoIIQ*-F-AscI/Term-R-EcoRI and cloned between the AscI and EcoRI restriction sites of pP*x+*’*gfp*’. This construction was used to detect GFP production upon xylose induction among the non-sporulating cells. | This study |
| p*PspoIIQ*’*mCherry*-P*ykuN2*’*gfpAAV* | The promoter region of the *ykuN2* gene was amplified by PCR from the chromosome of *B. thuringiensis* 407 using primer pairs Pykun2-F-SalI/Pykun2-R-AscI and cloned between the SalI and AscI restriction sites of pP*spoIIQ*’StabSD’*comGAmCherry*’TermLambda-P*nprA*’*gfpAAV*. This construction allows to determine the RNA-Seq target gene promoter activity among the non-sporulating cells. | This study |
| pP*spoIIQ*’*mCherry*’-P*isdE1*’*gfpAAV* | This plasmid was constructed as above with the promoter region of *isdE1* gene using primer pairs PisdE1-F-SalI/PisdE1-R-AscI | This study |
| pP*spoIIQ*’*mCherry*’-P*dhbA*’*gfpAAV* | Same process as the plasmid above with the promoter region of *dhbA* gene using primer pairs PdhbA-F-SalI/PdhbA-R-AscI | This study |
| pP*spoIIQ*’*mCherry*’-P*BTB_c10430*’*gfpAAV* | Same process as the plasmid above with the promoter region of *BTB_c10430 gene* using primer pairs PBTB_c10430-F-SalI/PBTB_c10430-R-AscI | This study |
| pP*spoIIQ*’*mCherry*’-P*katE*’*gfpAAV* | Same process as the plasmid above with the promoter region of *katE1* gene using primer pairs PkatE1-F-SalI/PkatE1-R-AscI | This study |
| pP*spoIIQ*’*mCherry*’-P*sodA1*’*gfpAAV* | Same process as the plasmid above with the promoter region of *sodA1* gene using primer pairs PsodA1-F-SalI/PsodA1-R-AscI | This study |

**Table S2. Strains used in this study**

| **Name** | **Relevant information** | **Référence** |
| --- | --- | --- |
| Bt (pHT304) | *B. thuringiensis* 407^-^ carrying the empty pHT304 vector and used as a fluorescence^-^ control. | [8] |
| Bt (pP*spoIIQ*’*mCherry*-P*nprA*’*gfpAAV*) | *B. thuringiensis* 407^-^ in which we measure the activity of the promoter of *nprA*, using a reporter gene encoding an unstable GFP, as well as the activity of the promoter of *spoIIQ*, using mCherry, to study the behavior of the Spo^-^ cells | This study |
| Bt (pP*spoIIQ*’*mCherry*’) | *B. thuringiensis* 407^-^ in which we measure the activity of the promoter of *spoIIQ*, using a reporter gene encoding mCherry to study the Spo^-^ cells, to use in combination with green molecular dyes. | This study |
| Bt (pP*x+*’*gfp*’-P*spoIIQ*’- *mCherry*) | *B. thuringiensis* 407^-^ in which we measure the activity of the promoter of *spoIIQ,* using a reporter gene encoding mCherry coupled to the transcriptional fusion between Pxyl+ and GFP to determine GFP synthesis ability of the Spo^-^ cells upon xylose induction. | This study |
| Bt (p*PspoIIQ*’*mCherry*-P*ykuN2*’*gfpAAV*) | *B. thuringiensis* 407^-^ in which we measure the activity of the promoter of *ykun2*, using a reporter gene encoding an unstable GFP, as well as the activity of the promoter of *spoIIQ*, using mCherry, to study the behavior of the Spo- cells. | This study |
| Bt (pP*spoIIQ*’*mCherry*’-P*isdE1*’*gfpAAV*) | Same as the strain above with the promoter region of *isdE1* gene. | This study |
| Bt (pP*spoIIQ*’*mCherry*’-P*dhbA*’*gfpAAV*) | Same as the strain above with the promoter region of *dhbA* gene. | This study |
| Bt (pP*spoIIQ*’*mCherry*’-P*BTB_c10430*’*gfpAAV*) | Same as the strain above with the promoter region of *BTB_c10430* gene. | This study |
| Bt (pP*spoIIQ*’*mCherry*’-P*katE1*’*gfpAAV*) | Same as the strain above with the promoter region of *katE1* gene. | This study |
| Bt (pP*spoIIQ*’*mCherry*’-P*sodA1*’*gfpAAV*) | Same as the strain above with the promoter region of *sodA1* gene. | This study |

**Table S3. Oligonucleotides used in this study (pink letters indicate restriction sites)**

| **Name** | **Sequence** |
| --- | --- |
| PnprA-F-XbaI | gctctagaGCCGGAAAGGGTTTTTTCAATATTTG |
| PnprA-R-AscI | tggcgcgccGCTTTCTTACCAGTCGCTCC |
| PspoIIQ-F-SphI | acatgcatgcGCATCTTCGGTTGAAGTTCTAC |
| PspoIIQ-R-XbaI | gctctagaCATCACCTCAGCAATCATTTTGAAC |
| Term-F-KpnI | ggggtaccGATCTCTGCAGTCGCGATGATTAATTAATTC |
| Term-R-EcoRI | ggaattcCGCAACGTTCTTGCCATTGCTGC |
| StabSD-F-XbaI | gctctagaTCTTGAAAGGAGGGATGCCTAAAAA |
| StabSD-R-BamHI | cgggatccATAAAATGATTTTTCATAAATCCA |
| comGAmC-F-BamHI | cgggatccTTAAGGAGGTGACACCATGAATGGG |
| mC-R-KpnI | ggggtaccTTACTTATATAATTCATCCATTCCAC |
| PspoIIQ-F-SalI | acgcgtcgacGCATCTTCGGTTGAAGTTCTAC |
| Term-R-SphI | acatgcatgcGCAACGTTCTTGCCATTGCTGC |
| Term-R-AscI-XbaI-EcoRI | ggaattccgctctagaggcgcgccGCAACGTTCTTGCCATTGCTGC |
| PspoIIQ-F-AscI | tggcgcgccGCATCTTCGGTTGAAGTTCTAC |
| Pykun2-F-SalI | acgcgtcgacAAAAAAGCACAGATGATTGTATAGT |
| Pykun2-R-AscI | tggcgcgccCATCTAAACTAACTTTAATTAAATC |
| PisdE1-F-SalI | acgcgtcgacTATTGTTAACTAGAGCGCGGCGAAA |
| PisdE1-R-AscI | tggcgcgccAGACGCTTTCTCGTCCCCTTTGGCA |
| PdhbA-F-SalI | acgcgtcgacTATACAAATCTTCTATAACACTATG |
| PdhbA-R-AscI | tggcgcgccCTAAAAACATTTTGGCAACAACACT |
| PBTB_c10430-F-SalI | acgcgtcgacTAAATGCAATTTGGCAACAAACTAA |
| PBTB_c10430-R-AscI | tggcgcgccCGTAAGCATAGTACTGCTTAAATAA |
| PkatE1-F-SalI | acgcgtcgacCGAAAAGAATTATCTTAAAAGCCAA |
| PkatE1-R-AscI | tggcgcgccCTTGGTTTGTTGTTAAAGCATGTTT |
| PsodA1-F-SalI | acgcgtcgacCATATCCATTTCGCATGTTTATTA |
| PsodA1-R-AscI | tggcgcgccGGATGTTCATTGTTTCTTTGTCCAA |

**LEGENDS**

**Table S4 – Differential gene expression for necrotrophism genes at 7 dpi**

Table of necrotrophism differential expression in insect cadavers at 7 dpi compared to exponential growth phase cells (left column), stationary growth phase cells (mid column) or in bacteria harvested in stationary growth phase compared to exponential growth phase (right column). Necrotrophism genes were retrieved necrotrophism regulon published by Dubois et al., 2016 [11].

Purple indicates low expression (log2FC ≤-2), blue indicates high expression (log2FC value ≥2), gray indicates an adjusted p-value>0,01 or -2 > Log2FC < 2

**Table S5 – Differential gene expression for sporulation genes at 7 dpi**

Table of sporulation differential expression insect cadavers at 7 dpi compared to exponential growth phase cells (left column), stationary growth phase cells (mid column) or in bacteria harvested in stationary growth phase compared to exponential growth phase (right column). Sporulation gene were retrieved from KEGG and subtiwiki database.

Purple indicates low expression (log2FC ≤-2), blue indicates high expression (log2FC value ≥2), gray indicates an adjusted p-value>0,01 or -2 > Log2FC < 2

**Table S6 – Differential gene expression for germination genes at 7 dpi**

Table of germination differential expression insect cadavers at 7 dpi compared to exponential growth phase cells (left column), stationary growth phase cells (mid column) or in bacteria harvested in stationary growth phase compared to exponential growth phase (right column). Sporulation gene were retrieved from KEGG and subtiwiki database.

Purple indicates low expression (log2FC ≤-2), blue indicates high expression (log2FC value ≥2), gray indicates an adjusted p-value>0,01 or -2 > Log2FC < 2
